# Insights into mammalian TE diversity via the curation of 248 mammalian genome assemblies

**DOI:** 10.1101/2022.12.28.522108

**Authors:** Austin B. Osmanski, Nicole S. Paulat, Jenny Korstian, Jenna R. Grimshaw, Michaela Halsey, Kevin A.M. Sullivan, Diana D. Moreno-Santillán, Claudia Crookshanks, Jacquelyn Roberts, Carlos Garcia, Matthew G. Johnson, Llewellyn D. Densmore, Richard D. Stevens, Zoonomia Consortium, Jeb Rosen, Jessica M. Storer, Robert Hubley, Arian F.A. Smit, Liliana M. Dávalos, Kerstin Lindblad-Toh, Elinor K. Karlsson, David A. Ray

**Affiliations:** Department of Biological Sciences, Texas Tech University, Lubbock, TX, USA; Department of Natural Resources Management and Natural Science Research Laboratory, Museum of Texas Tech University, Lubbock, TX 79409, USA; Institute for Systems Biology, Seattle, WA, USA; Department of Ecology & Evolution, Stony Brook University, Stony Brook, NY, USA; Consortium for Inter-Disciplinary Environmental Research, Stony Brook University; Stony Brook, NY, USA; Department of Medical Biochemistry and Microbiology, Science for Life Laboratory, Uppsala University; Uppsala, 751 32, Sweden; Broad Institute of MIT and Harvard; Cambridge, MA 02139, USA; Program in Bioinformatics and Integrative Biology, UMass Chan Medical School; Worcester, MA 01605, USA; Program in Molecular Medicine, UMass Chan Medical School; Worcester, MA 01605, USA

## Abstract

We examined transposable element (TE) content of 248 placental mammal genome assemblies, the largest *de novo* TE curation effort in eukaryotes to date. We find that while mammals resemble one another in total TE content and diversity, they show substantial differences with regard to recent TE accumulation. This includes multiple recent expansion and quiescence events across the mammalian tree. Young TEs, particularly LINEs, drive increases in genome size while DNA transposons are associated with smaller genomes. Mammals tend to accumulate only a few types of TE at any given time, with one TE type dominating. We also found association between dietary habit and the presence of DNA transposon invasions. These detailed annotations will serve as a benchmark for future comparative TE analyses among placental mammals.

**One-Sentence Summary:** A *de novo* assessment of TE content in 248 mammals finds informative trends in mammalian genome evolution.

## Main Text

Barbara McClintock became a scientific pioneer in the field of genomics with her Nobel Prize winning discovery of transposable elements (TEs), DNA sequences that can mobilize themselves in host genomes (*1*). A ubiquitous component of nearly all eukaryotes (*2*), TEs are typically classified into two major groups based on their mobilization mechanism (*3*). Class I elements, also known as retrotransposons, utilize an RNA intermediate during transposition allowing replication throughout the genome in a copy-&-paste style of mobility (*4*). Class I elements can be sorted further into three subcategories: Short INterspersed Elements (SINEs), Long INterspersed Elements (LINEs), and LTR-retrotransposons (*5*). SINEs are non-autonomous elements and depend on the presence of functional LINE elements, which contain anywhere from 1-3 open reading frames encoding the necessary proteins for mobilization. Class II elements, also known as DNA transposons, employ a DNA intermediate and can also be subdivided. TIR-like DNA transposons such as hATs, piggyBacs, and TcMariner transposons utilize a cut-&-paste mechanism by using transposase enzymes to catalyze the TE’s relocation (*6*). Helitrons, a second subcategory of Class II elements utilize a rolling-circle mechanism (*7*). The final subcategory of known DNA transposons are Maverick elements, which are thought to be derived from viruses as they have homologous genes coding for DNA polymerase and retroviral-like integrase (*8*).

An increase in activity from either class of elements can lead to drastic alterations in genome architecture (*9*). A variety of changes, including insertions, duplications, translocations, deletions, and inversions can result from TE mobilization and accumulation (*9*). For instance, the *AMAC1* (acyl-malonyl condensing enzyme 1) gene, coding for a protein essential for breaking down phytanic acid from meat and dairy foods, has undergone multiple recent gene duplications mediated by SVA retrotransposons in the human genome (*10, 11*). In addition to these structural variants, the proliferative mechanisms of TE mobilization tend to cause eukaryotic genome sizes to linearly correlate with TE abundance (*2*).

Increasing evidence indicates that TE-derived sequences have substantially influenced the evolutionary histories of the organisms they occupy, even contributing to major evolutionary innovations benefitting host organisms. Examples include recent TE insertions into genes involved with insecticide resistance of the cotton bollworm (*12*); the rapid adaptation leading to melanistic phenotypes of peppered moths in the soot ridden environment of British industrialization (*13*); and the myriad of endogenous retroviruses that have contributed novel regulatory functions to the development and evolution of the mammalian placenta (*9, 14*). The overwhelming majority of TE insertions, however, result in selectively neutral alterations in genome architecture, often showing no perceptible effect on host fitness (*15*). That being said, deleterious insertions occur and impairments in gene function are possible outcomes of TE mobilization which can lead to a wide variety of genetic diseases (*9*).

As a result, numerous genomic TE defense mechanisms have evolved to combat TE activity by either regulating TE transcription or by targeting their intermediates to prevent integration into the genome (*3*). These defense mechanisms explain, in part and in some organisms, why few TE families retain the ability to mobilize over long periods of evolutionary time (*16*). For example, among the ∼868,000 L1 insertions in the human genome, few are thought to be retrotransposition-competent and many of these exhibit cell-type specific mobilization profiles (*3, 17*). Alternative to or in conjunction with the aforementioned scenario of low numbers of functionally mobile TEs among some categories of elements, genomic drift and the corresponding effects of fixation events among bottlenecked populations gives rise to another explanation for varying levels of TE accumulation in different genome assemblies (*18*).

All these facets suggest that determining TE dynamics is key to understanding how genomes evolve and function. Thus, TE curation and annotation is one of the most important initial investigative steps in any description of a *de novo* genome assembly. Unfortunately, this step is often relegated to an afterthought rather than performing a time-intensive *de novo* TE curation effort (*19*). As a result, many genome assemblies are misunderstood from a TE perspective (*19*). As the scientific community improves genome sequencing and assembly, the lack of thorough and accurate TE annotation promises to become a major problem, especially in the face of the number of large-scale genome sequencing initiatives now underway (*20-24*).

The Zoonomia project, described in (*24*), represents an opportunity to gain substantial knowledge of the diversity of TEs in an important vertebrate clade, Mammalia. Here, we fill this knowledge gap by providing complete, *de novo* TE annotations of 248 Zoonomia mammalian genome assemblies using homology, *de novo*, and manual annotation approaches.

## Results

### General TE trends among mammals

RepeatModeler (*25*), a *de novo* TE discovery tools, was used to examine 248 mammalian genome assemblies yielding 25,025 putative TE starting queries. After initial curation and elimination of duplicates, an iterative curation process consisting of between 1 and 19 rounds of detailed curation (*19*), depending on the species (see Methods), yielded a library consisting of 8,263 novel consensus sequences. That library was combined with known TEs to create a comprehensive mammalian TE library. This library, consisting of 25,676 consensus sequences, was used to mask all assemblies. The dynamics of TE biology and intricacies of TE detection lend themselves to a degree of false detection. For example, some TE families are chimeras of multiple elements or they may contain similar core sequence components. To evaluate the potential for false positives, we took advantage of an idiosyncrasy of TE biology in bats. A family of bats, the Vespertilionidae, is, to our knowledge the sole mammalian family to have incorporated a type of rolling circle transposon, Helitrons, into their TE repertoire (*3*). True Helitrons in mammals have not been detected outside of Vespertilionidae. Thus, any Helitrons detected outside of vesper bats, would likely be a false positive. RepeatMasker (*26*) detected Helitrons in non-vesper mammals at a rate of 0.0013 ± 0.0019, suggesting a low false positive rate.

Previous work has suggested that the largest single classifiable component of a typical mammalian genome is TEs (*27*) and our data (Fig. 1) corroborate this. As noted previously by Elliott & Gregory in 2015 (*2*), genome size linearly correlates with the percentage of TE content within a genome and this is again supported (Fig. 1, Table S1). Overall, TE content in each of the examined species ranges from a low of 27.6% in the star-nosed mole (*Condylura cristata*) to 74.5% in the aardvark (*Orycteropus afer*) (Table S2, Fig. 1), with a distinct tendency to cluster in the middle of that range (average TE proportion: 45.6%, average genome size: 2.67Gb). The hazel dormouse (*Muscardinus avellanarius*) and Brazilian guinea pig (*Cavia aperea*) represent the extremes of this middle cluster with 65.8% and 28.1% total TE content, respectively.

**Fig. 1.**
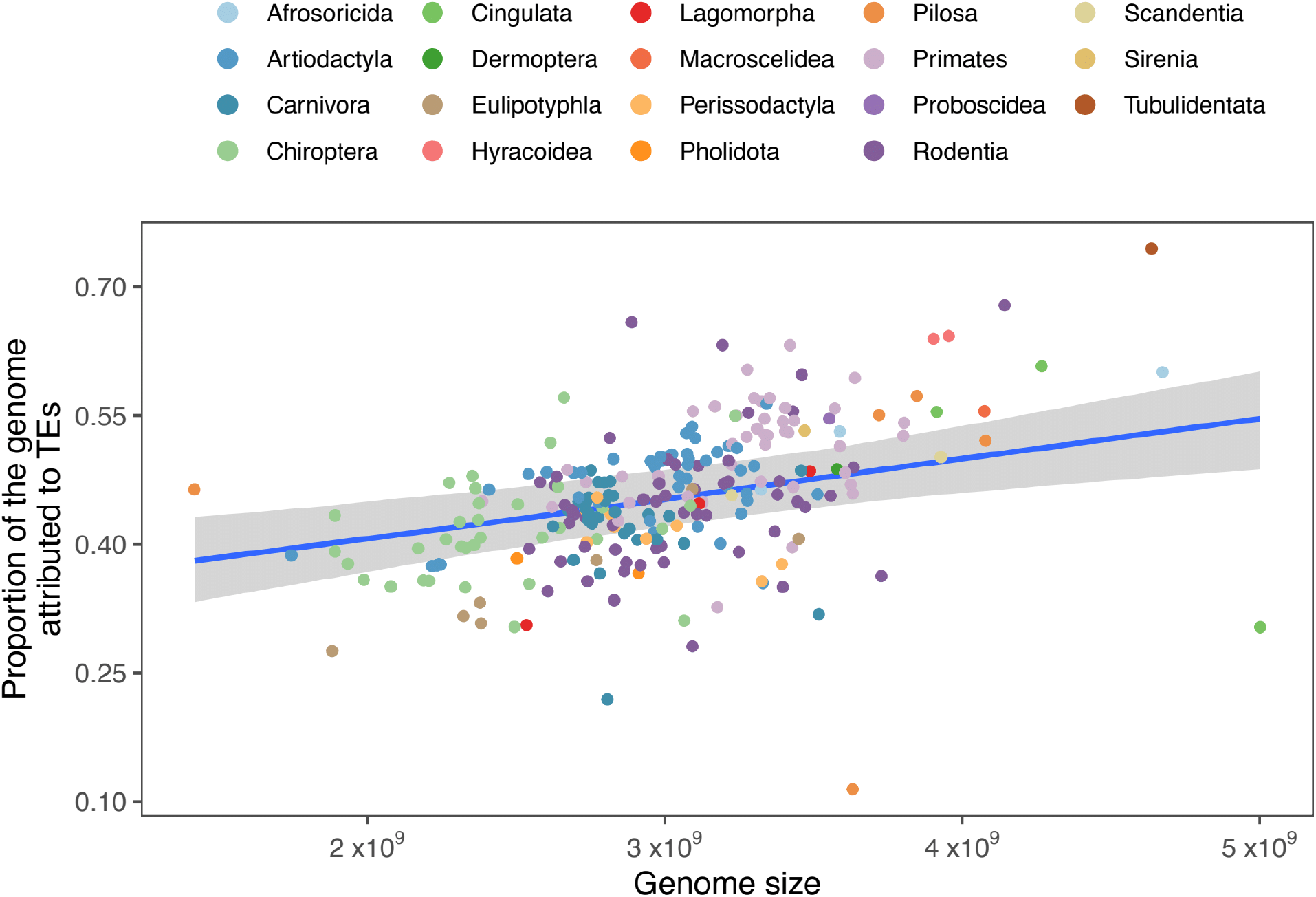
Correlation of total genomic TE content and the size, in base pairs, of the genome. Due to the log transformation and scaling of assembly size for the hierarchical Bayesian analysis, and resulting back-transformation, the x-axis values are approximately rendered. Blue line indicates the line of best-fit and shaded area is the 95% high probability density of the fit. The r^2^ for this relationship was estimated at 0.54 (95% high probability density 0.42, 0.64).

Assembly quality may impact the accuracy of TE annotation, but we could find no statistically significant trend among taxa. For example, lower quality assemblies as measured by N50 or BUSCO completeness did not yield lower or higher rates of observed TE accumulation (Fig. S1 and S2).

### TE variation among mammals

When examining TE content from all categories across the mammalian tree, we find some general trends. For example, SINEs and LTR retrotransposons are more prevalent in Euarchontoglires while LINEs dominate most other lineages, especially the bovids (Fig. 2). However, we find placental mammals are generally similar with regard to overall TE proportions, reflecting the tendency to retain older insertions that occurred in the common ancestor of mammals. LINEs and SINEs always make up most TE abundance both in copy number and in total genomic percentage. LINEs occupy between 8.2% and 52.8% of the genomes examined, averaging 22.6%. SINEs occupy on average 10.5% of the mammalian genome (range 0.4%-32.1%) (Table S3) while LTR retrotransposons, DNA transposons, and rolling-circle transposons (RC) are substantially rarer; 7.8% (range 2.0%-17.8%), 3.5% (range 0.5%-8.4%), and 0.5% (range 0.01%-19.7%), respectively.

**Fig. 2.**
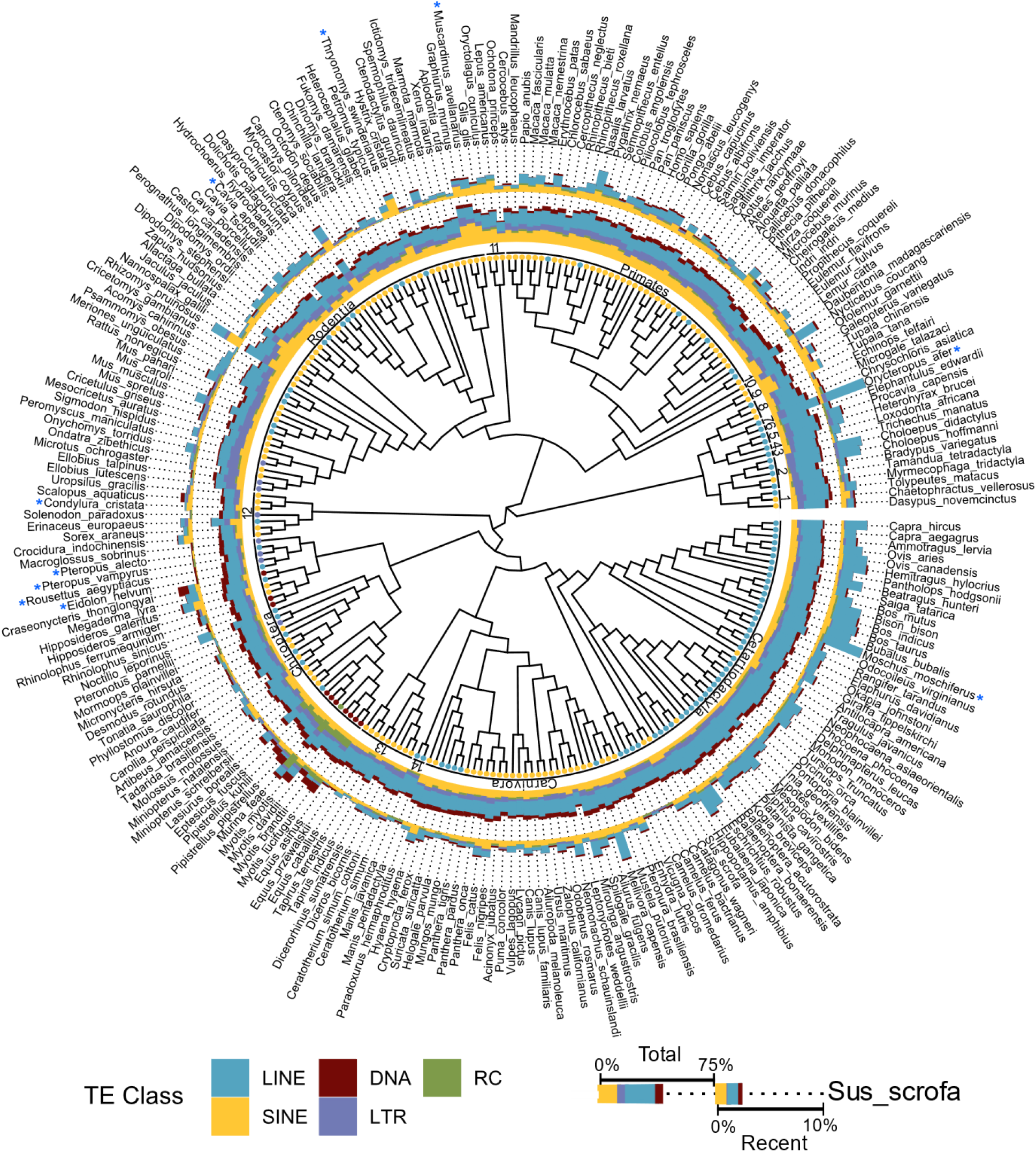
Total and young TE genomic proportions by species within a phylogenetic context. Dots at branch tips indicate the TE class most prevalent among recent TE insertions (insertions with <4% divergence from the relevant consensus TE). The ring immediately following the branch tip dots indicates the mammalian order for reach respective species. Orders represented by numbers include: 1) Cingulata, 2) Pilosa, 3) Sirenia, 4) Proboscidea, 5) Hyracoidea, 6) Macroscelidea, 7) Tubulidentata, 8) Afrosoricida, 9) Scandentia, 10) Dermoptera, 11) Lagomorpha, 12) Eulipotyphla, 13) Perissodactyla, 14) Pholidota. The inner ring of stacked-bar data depicts the total percentage of the genome attributed to the five main categories of TEs: DNA transposons, LINEs, SINEs, LTRs, & Helitrons. The outer ring of stacked-bar data shows the percentage of the genome derived from recently inserted TEs. Cladogram adapted from (*65*).

Examination of younger insertions, those with divergences averaging <4% from their respective consensus, provides a picture of these genomes that is more dynamic, revealing substantial differences in accumulation from each category of TE (Table S4). Some lineages, such as the pteropodid bats (*Pteropus alecto, P. vampyrus, Eidolon helvum*, and *Rousettus aegyptiacus* in Fig. 2), exhibit essentially no recent accumulation by any TE category while others have experienced massive expansions in one or more categories. The aardvark (*Orycteropus afer*) and musk deer (*Moschus moschus*) for instance, show substantial LINE accumulation over the past ∼20 million years.

To examine these trends more closely, we conducted a redundancy analysis (RDA) for both orders and families to identify the major axes of variation in TE composition that were related to either order or family affiliation of taxa (Fig. 3). This analysis suggests a strong phylogenetic component to variation in TE composition among clades at the levels of order and family. Eleven orders of mammals were significantly correlated with at least one of the two axes and these orders were quite variable in terms of association with different TE types. The first two major axes of variation in TE accumulation in analyses examining orders accounted for approximately 27.2% of the variation and this was highly significant (P < 0.001). The first major axis was positively related to the number of young TEs generally, and to young LINEs, LTRs, and SINEs, which are all obligately replicative. Unsurprisingly given this characteristic, genome size was also positively correlated with this axis. This axis was negatively related to young DNA transposons and young rolling circle transposons. The second major axis of TE composition related to ordinal affiliation was positively related to the number of young DNA transposons, rolling circle transposons, LINEs and young TEs more generally, but negatively related to young LTRs, SINEs, and to genome size.

**Fig. 3.**
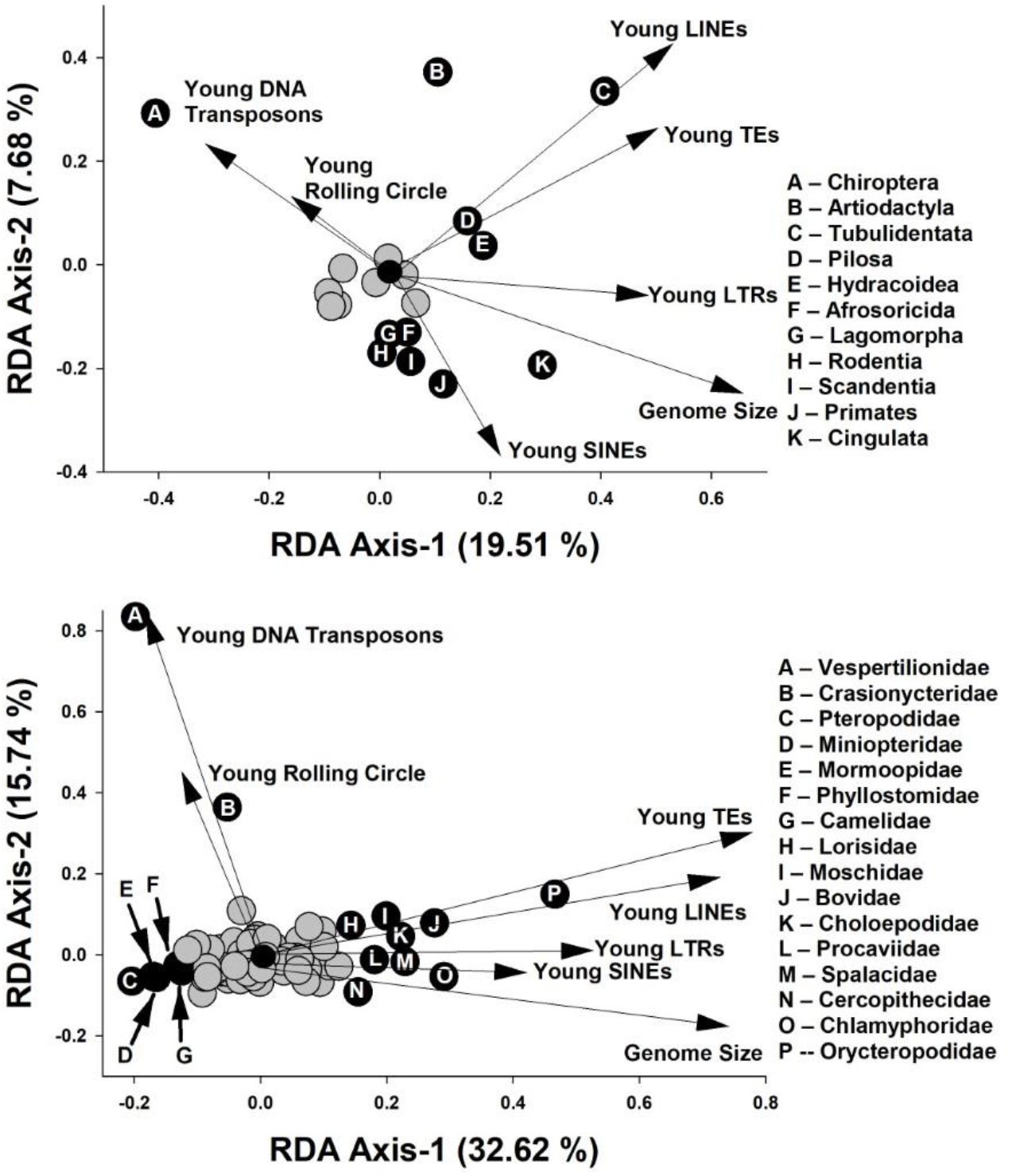
Redundancy analyses examining major axes of variation in TE accumulation and genome size related to orders (above) and families (below) of mammals. Arrows represent significant correlations TE types with the first two RDA axes. Each axis reflects changes in TE composition related to ordinal (above) or familial (below) affiliation of taxa used in analyses. Gray circles represent orders or families that were not significantly correlated to at least one of the RDA axes whereas black circles represent orders or families with significant correlations.

Similar associations are seen at the family level. Families of mammals accounted for approximately 49.9% of variation in TE composition, and this was highly significant (Fig. 3; P < 0.001). As with orders, the first major axis of variation was positively related to the same categories of TE and to genome size. Correlations of young DNA transposons and young rolling circle TEs were weaker than for orders, likely due to the lineage specificity of those element types (see below), while positive associations of all other TE types were stronger. The second major axis was positively related to the number of young DNA transposons, rolling circle transposons, LINEs, and young TEs generally and negatively related to genome size. Fourteen families of mammals were significantly correlated with at least one of these two axes and these families were variable in terms of association with different TE types.

### TE diversity

An increasingly useful avenue of inquiry among whole-genome TE analyses draws from community ecology (*28*). Of interest is the application of community diversity measures rendered on a genomic scale (*29*). We followed these lines of inquiry by investigating the diversity of recent TEs in each genome by calculating two diversity indices and applying them to our data, Shannon diversity index (*30*) and Pielou’s *J* (*31*).

Shannon diversity is a measure of overall diversity in a population of objects while Pielou’s *J* measures evenness by incorporating the relative numbers of each object, in this case, TE types (Table S5). Species with the highest diversity values include bats and rodents. Bat TE diversity was driven primarily by recent expansion of DNA transposons among Craseonycteridae, Vespertilionidae, Hipposideridae, Rhinolophidae, and Mollossidae and recent accumulation of both DNA transposons and rolling circle transposons in Vespertilionidae (Fig. 4).

**Fig. 4.**
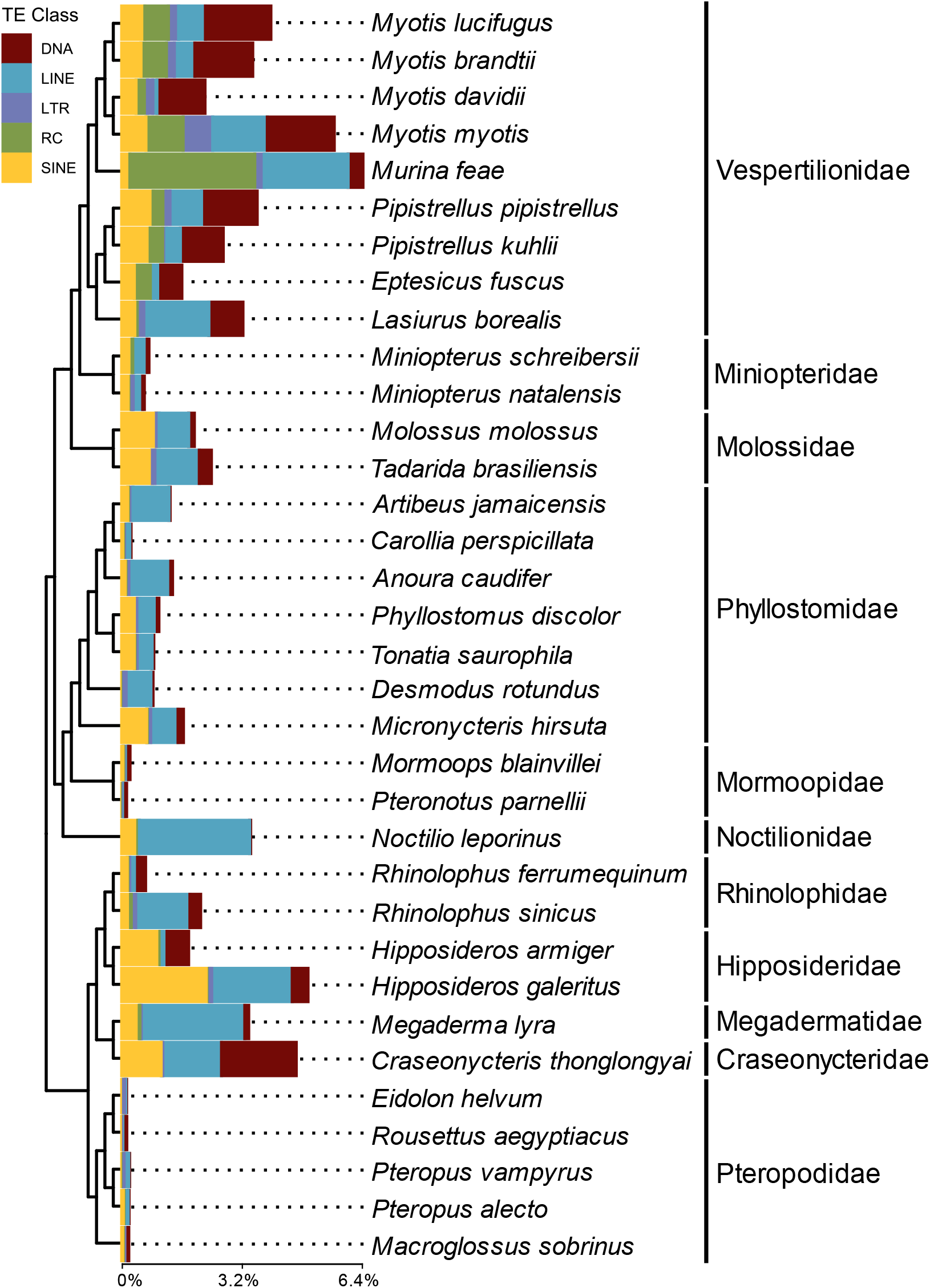
Stacked bar charts depicting proportions of recently accumulated TEs (<4% kimura from consensus TE) in bats. Data is organized by TE classification and plotted onto the tips of the chiropteran portion of the mammalian tree, adapted from (*65*).

In rodents, higher diversity among recently inserted TEs was driven by accumulations in LTR retrotransposons, which made up 10-53% of recent TE accumulation. The highest rate of recent LTR accumulation among the rodents was seen in members of Cricetidae and *Cricetomys gambianus*.

To investigate general trends in diversity index values in relation to TE accumulation patterns, we plotted values from recently deposited TEs vs. each diversity index (Fig. 5). Hierarchical Bayesian analyses indicate that both Shannon diversity and Pielou’s *J* exhibit significant negative relationships with increasing recent TE content; Shannon *H* (Fig. 5, Table S6) and Pielou’s *J* (Fig. 5, Table S7, Fig. S3). Thus, the downward trend in Pielou’s *J* suggests that mammalian genomes tend to accumulate individual TE types at any given period rather than multiple TE types accumulating simultaneously. This is exemplified in the aardvark, where LINEs are currently dominating the recently active mobilome while SINEs are the major recent contributor to the greater cane rat (*Thryonomys swinderianus*) genome (Fig. 2). However, clades of bats with recent DNA accumulation tend to refute this pattern.

**Fig. 5.**
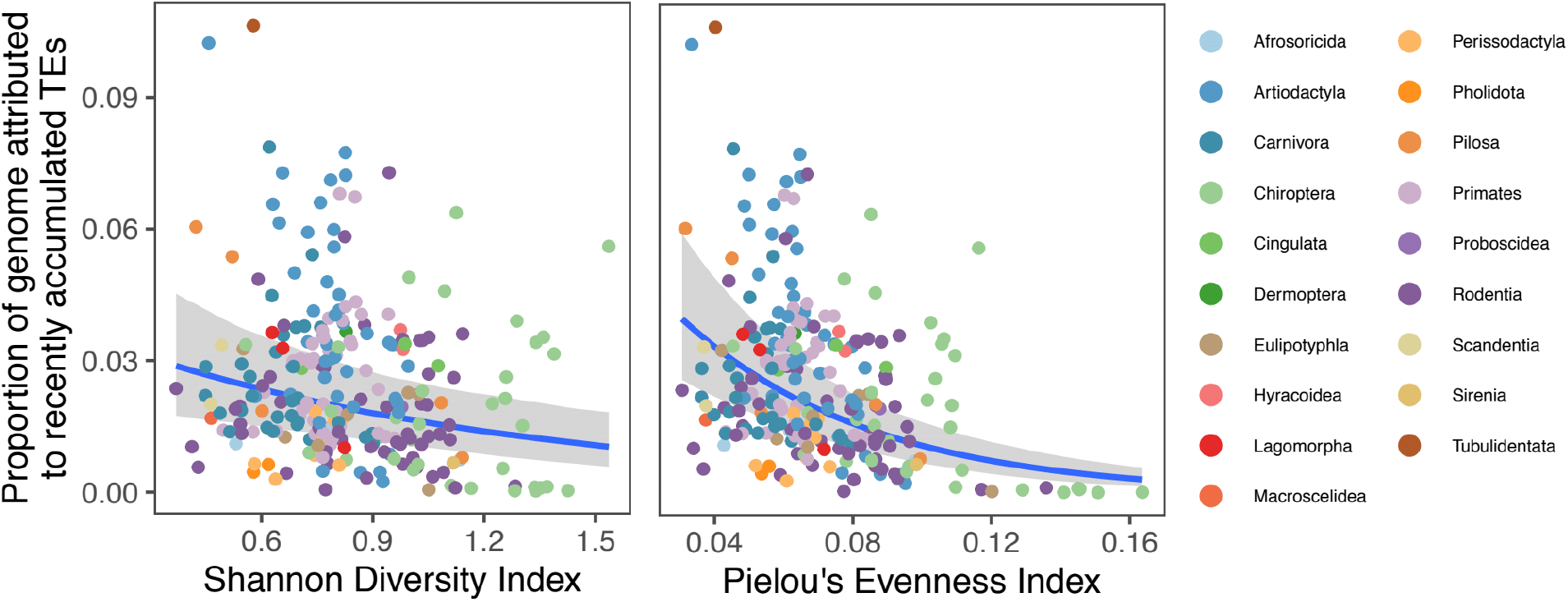
Recent mammalian TE diversity in relation to Shannon *H* (left) and Pielou’s *J* (right). Blue line indicates the line of best-fit and shaded area is the 95% high probability density of the fit. The r^2^ for *H* was estimated at 0.67 (95% high probability density 0.52, 0.78), and for *J* it was 0.69 (95% high probability density 0.56, 0.79).

### DNA transposons and diet

The lineage specificity of the DNA transposon diversity described above suggests horizontal transfer (HT) as a potential source of novel TE invasions in certain mammalian genomes. To investigate patterns that may explain how such HT events may occur, we examined the potential for life history to play a role. We hypothesized that differences in diet may allow select species to come in contact with vectors for TEs (*14, 32*), which increase the likelihood of successful invasion of mammalian genomes. DNA transposon-rich food sources such as many arthropods, and non-mammalian vertebrates may offer greater potential for HT to some species compared to those that eat plants. Hierarchical Bayesian analyses indicate that carnivorous mammals tend to accumulate more recent DNA transposons in their genomes than non-carnivores (Fig. 6A; Table S8). This pattern is best exemplified in the cetartiodactyls (Fig. 6B). Recent DNA transposon accumulation is seen on average 20x more among the cetaceans than other artiodactyls. Carnivorous bats, however, did not have statistically higher accumulations of recent DNA transposons than herbivorous bats (Fig. 6C). Our datasets of primates and rodents did not reveal any statistical difference of recent DNA transposon accumulation between herbivores and omnivores (Fig. 6D-E).

**Fig 6.**
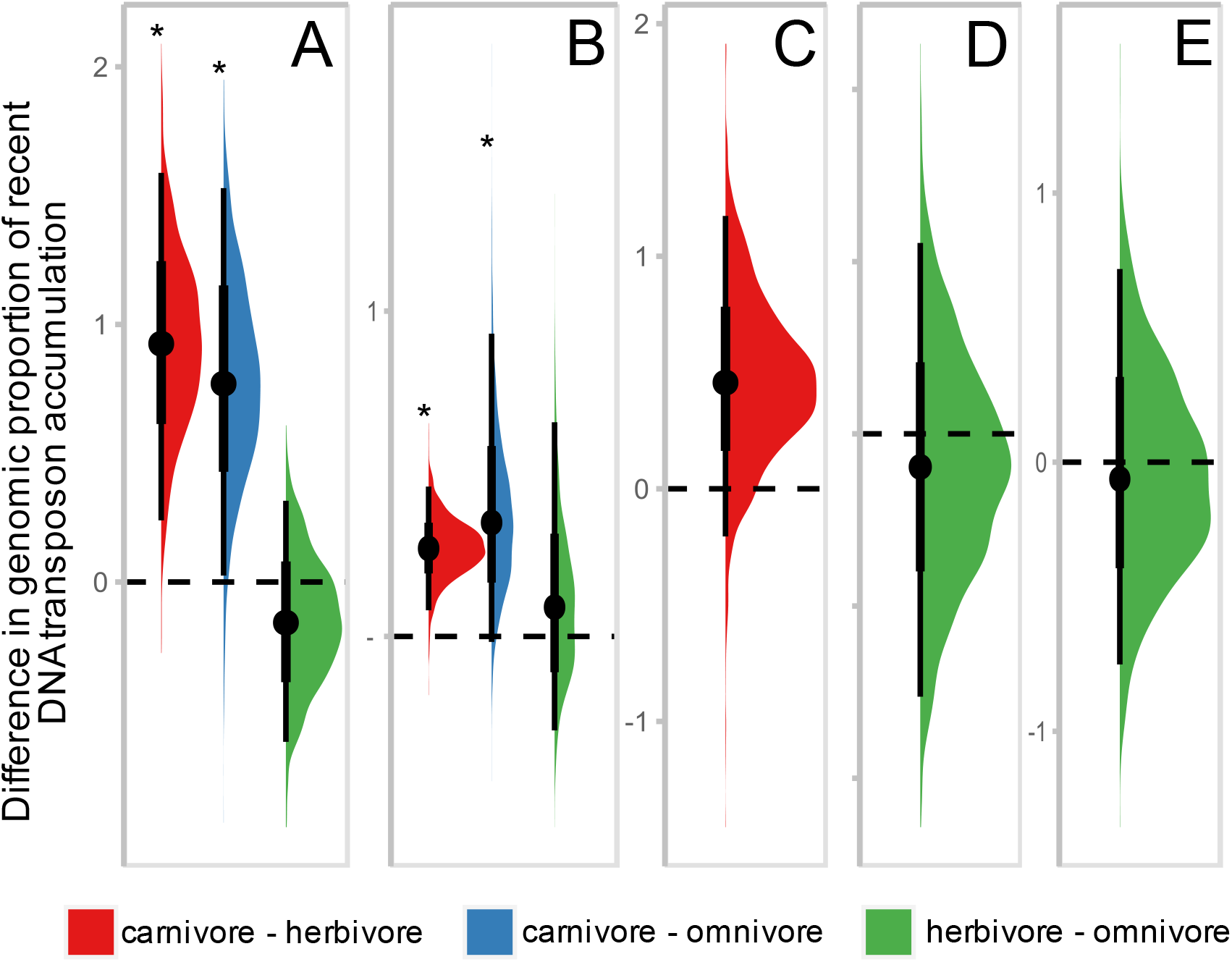
Half eye plots depicting fold differences in recent DNA transposon accumulation among three dietary phenotypes: carnivore, herbivore, and omnivore. Instead of showing the estimated values for each of the diets, these plots depict the fold ratio between each diet pair, so that the plot itself shows statistical significance. Comparisons for which the thin line does not overlap with 1 are significant, indicated by *. Plots correspond to the following taxonomic groups: A) placental mammals (r^2^ estimated at 0.92 (95% high probability density 0.79, 0.97)), B) Artiodactyla (r^2^ estimated at 0.64 (95% high probability density 0.32, 0.78)), C) Chiroptera (r^2^ estimated at 0.34 (95% high probability density 0.02, 0.86)), D) Primates (r^2^ estimated at 0.18 (95% high probability density 0.00, 0.58)), E) Rodentia (r^2^ estimated at 0.07 (95% high probability density 0.00, 0.28)).

## Discussion

As our ability to generate high quality genome assemblies in rapid succession improves, the need to curate TEs in those assemblies will only increase. Toward that end, we performed a *de novo* assessment of the TE content of 248 mammal genome assemblies in what is, to our knowledge, the largest comprehensive TE curation effort to date. This represents an increase of ∼58% compared to known mammalian TEs in RepBase as of 2019, when we began. Given the numerous impacts that TEs are known to have at multiple levels of genome organization and function, this increased knowledge will serve as a particularly valuable resource for anyone interested in mammalian genomics and evolution. The full set of TE consensus sequences is available for download from the Dfam (*33*) database.

Previous work has noted that genome size among mammals is relatively constrained (*34*) and this work does not contradict that observation. Despite this constraint, our effort notes that there is substantial variation in rates of accumulation in the recent mammalian past. Indeed, we found that there is substantial diversity in TE accumulation patterns among mammals, suggesting distinct TE-induced pressures on those genomes over evolutionary time and, likely, distinct differences in the ability of eutherians to defend their genomes against TEs. These differences represent an excellent opportunity for future researchers to investigate how TE defenses evolve and respond to differing TE loads.

Another avenue of such research is to further investigate TE accumulation through the lens of ecology and environment, an idea that has been discussed previously (*14*). Our data demonstrate that carnivorous lineages tend to harbor an excess of recently accumulated DNA transposons when compared to herbivorous taxa. The tendency of meat-eating mammals to have more recent DNA transposon accumulation as compared to their non-carnivorous counterparts suggests diet may play a significant role in a genome’s likelihood of experiencing HT from Class II TEs. This scenario is supported in part by a recent analysis of HT in predator-prey pairs and their shared parasites (*32*). Nevertheless, this finding is not uniform across mammalian orders and those varying patterns may reflect defenses against TE invasion (*3*), less availability of TEs in order-specific dietary items, or some combination of both.

Investigating mammalian TEs through the ecological lens also suggests that single TE types tend to dominate the mobilome during any given period (Fig. 5). This scenario is consistent with our current understanding of TE defense mechanisms. The current model of PIWI-mediated TE defense suggests that a new TE may invade or arise in a genome and enjoy a period of relatively unfettered mobilization. Eventually, the piRNA defenses generate an effective response and dampen the new TE’s impacts (*16, 35, 36*).

With regard to the prevalence of HT of DNA transposons in carnivores, our data support the hypothesis that the prevalence of HT of DNA transposons may be a consequence of the similar cellular environments of predator and prey and their necessarily shared environments and frequent interactions. Recent research has demonstrated the role that viruses and blood-feeding arthropods play in facilitating HT (*14, 32*). Frequent interactions would further facilitate HT by bringing such vectors into contact with both predator and prey. The similar cellular environments among animals (as opposed to mammals with plant-based diets) would further encourage the ready transfer of DNA transposons, which are already more amenable to HT due to their relatively weak dependence on a host’s cellular machinery to mobilize (*37*).

In conclusion, the annotation data provided here is essential for answering future questions related to emerging hypotheses around speciation such as the TE-Thrust Hypothesis, the Epi-Transposon Hypotheses, or the Carrier SubPopulation Hypothesis (*3, 38*). As anthropogenic change exacerbates the decline in effective population size for many of the species in our dataset, transposable elements might be the reservoir of genomic mutagens that future populations or species rely on.

## Materials and Methods

### Generating the mammalian TE library

A total of 248 genome assemblies of placental mammals were initially presented for analysis (table S2). For six species, higher quality assemblies were available via Bat1k, a similar, large scale genome sequencing and assembly effort (*21*). In those cases, we replaced the Zoonomia assembly with the higher quality version. Some assemblies were not used in the development of our final mammalian TE library due to one or more of the following reasons: 1) the assembly exhibited a low N50 value (<20,000) resulting in short contigs which are unsuitable for identifying longer TEs, 2) multiple artifacts of assembly error were observed at TE sites which yielded implausible consensus sequences, 3) a thorough species-specific TE annotation had already been performed and is available from RepBase (Genetic Information Research Institute) (*39*), previous work from our own laboratory, or work conducted by a collaborator. This left us with 205 species as substrates for TE curation (table S2).

Mammalian genomes have only a minimal tendency to remove older TE insertions from the genome (*40*). Thus, the majority of older TE families that mobilized in the common ancestor or early in the mammalian diversification were likely already characterized through efforts that focused on any of several model organisms such as human, mouse, rat, pig, dog, cat, and horse (*41-47*). To avoid wasted effort on re-curation of these shared and previously described TEs, we focused our manual curation efforts on identifying newer putative TEs that underwent relatively recent accumulation. We defined such ‘young’ insertions as TEs with sequences with K2P genetic distances less than 4% when compared to their respective consensus. For temporal orientation, a kimura divergence of 4% approximates 20mya or less since insertion, based on a general mammalian neutral mutation rate of 2.2×10^−9^ (*48*). The use of a general mutation rate allowed for consistency among K2P values in analyses, however it limits the accuracy of species-specific temporal estimations due to varying neutral mutation rates among placental mammals. Thus, results with divergence values of less than 4% are considered “young” and do not provide exact dates. This approach yielded mostly lineage specific TEs, many of which were yet to be described but some previously identified and shared elements were occasionally encountered (i.e. the Tigger family of Tc Mariner transpsosons and others), suggesting that we did not miss older but unidentified elements. Custom scripts associated with the identification of younger elements are available on zenodo (*49*).

For details of the curation process, see previous work from (*19*). Briefly, for each iteration of manual TE curation, new consensus sequences were generated from the 50 BLAST hits which shared the highest sequence identity to the consensus used in our BLAST query for that iteration. Custom pipelines accomplished this by aligning BLAST hits with MUSCLE (*50*), trimming alignments with trimAl (-gt 0.6 - cons 60) (*51*), and estimating a consensus sequence with EBMOSS (cons -plurality 3 -identity 3) (*52*). Files which resulted in fewer than 10 BLAST hits were discarded. To consider a consensus sequence ‘complete,’ the alignment needed to exhibit a pattern of random sequence at both the 5’ and 3’ ends, or after extension to a length of 7kb or greater, whichever came first.

Because the ubiquitous LINE-1 can introduce copies of any transcript into the genome, mammalian genomes have an unusually high number of processed pseudogenes (*53-55*). Including these in a repeat database would result in annotation of functional genes as TE copies. Comparisons with protein (domain) databases (https://www.ncbi.nlm.nih.gov/protein/, https://useast.ensembl.org/index.html) we found and removed 152 such entries, most characterized by a poly A tail. Small structural RNAs often occur in higher copy numbers partially because they are also substrates of LINE1 (*56*), and a further 49 entries were dismissed as models created from their genes and pseudogenes.

Two or three copies of interspersed repeats with very high copy numbers, usually but not exclusively SINEs, can often be found in tandem clusters. This occurs more than by chance do to target site preferences. For example, LINE-1 dependent SINEs insert in A-rich DNA, and such sites are introduced by their own poly A tails (*57*). These artifacts are often identified by de novo repeat finders but can be recognized when studying the seed alignments. Models will also have been built for the individual units and many copies will end at the joining region between the units, the joining region is more variable than the rest of the model. Over 210 models were such artifacts and eliminated.

Because in mammals the majority of LTR elements are represented by solo LTRs (*58*), Dfam (*33*) and Repbase (*39*) harbor separate models for the LTRs and the internal sequences. De novo repeat-finders like RepeatModeler often produce full elements or reconstruct a (partial) LTR and a fragment of the internal sequence. We split these models into their components, based on homology to well-defined LTRs and the presence of tRNA primer binding sites.

The combined original library contained several redundant models. Recognizing that models represent (fragments of) the same TE is complicated by incorrect base calls, indels, overextension, and incompleteness of the reconstruction as well as by the evolution of class I TEs in the genome: copies created at different evolutionary times or from different descendants of the ancestral TE (sometimes subtly) differ. A solid test for redundancy is to match the genome to all related models simultaneously and find that some models are always outcompeted by others or that models converge to the same consensus sequence. This could only be accomplished once the database was finalized, so we applied arbitrary but informed cutoffs. Before comparison to each other the low complexity tails of SINEs and LINEs were set to a standard length and short overextensions were trimmed, based on the expected signatures of terminal bases or target site duplications. Differences between models at possible (highly mutagenic) CpG sites were ignored. Dependent on class and age, elements were removed with alignment scores against another model with a more complete sequence or a better seed alignment that were between 90-95% of the score against itself. Partially overlapping fragments of potentially the same TE were not addressed at this point.

We eliminated duplicated entries only when they were built from the same assembly. The same TE can be reconstructed from the genomes of different species if it was active before their speciation time, but with our current approach we could not estimate if a repeat was shared or lineage-specific and merely similar. Thus, in Dfam (*33*) each of the models of this study currently is associated with only one species and will not be matched when a same model is present in another species library.

To confirm the TE type, each sequence in the library was subjected to a custom pipeline (*49*) which used: blastx to confirm the presence of known ORFs in autonomous elements, RepBase (*39*) to identify known elements, and TEclass (*59*) to predict the TE type. We also used structural criteria for categorizing TEs. DNA transposons were identified as elements with visible terminal inverted repeats. Rolling circle transposons were required to have identifiable ACTAG at one end. Putative SINEs were inspected for a repetitive tail as well as A and B boxes. SINEs also were classified by comparison to a database of SINE modules(*33*): 800 small RNA class III promoter regions, 150 core regions and 5500 3’ ends of LINE elements (which SINEs often share). LTR retrotransposons and solo LTRs were required to have recognizable hallmarks, such as: TG, TGT, or TGTT at their 5’ and the inverse at the 3’ ends, and the presence of a polyadenylation signal. LTR classes could often be assigned by (indirect) sequence homology to a coding internal sequence, when present. After this process, 8263 models and their seed alignments were submitted to Dfam (*33*).

Once the final mammalian TE library was created, we used RepeatMasker-4.1.0 to mask the genome assemblies. Postprocessing of output was performed using the rm2bed.py utility included with RepeatMasker, which merges overlapping hits and converts the output to bed format.

### Plotting TE variation using ordination

To characterize the major axes of variation of young TE accumulation among taxa we conducted a redundancy analysis for both orders and families. In these analyses, the number of base pairs attributed to of each TE Type as well as genome size for each taxon (order or family) was the dependent matrix and dummy variables (*60*) assigning a species to either family or order was the independent matrix. Redundancy is a multivariate regression that aims to examine the amount of variation and its statistical significance in the dependent matrix that can be accounted for by the independent matrix. Associations among variables where quantified based on a correlation matrix and significance was determined based on 9999 permutations of the original datasets. Redundancy analyses were performed in Canoco version 5 (*61*).

### Test for association between TE proportions and assembly size, two diversity indices, and diets

The three objectives of these analyses included: 1) quantifying the association, if any, between the total TE proportion in genome and assembly size; 2) estimating the difference in proportions of recently accumulated DNA transposons within a genome among species with different diets; 3) and quantifing the association, if any, between recent TE proportion in a genome and two diversity indices.

### Diversity indices

An increasingly useful avenue for characterizing TE accumulation draws on community ecology (*28*). Of particular interest is the application of community diversity measures rendered on a genomic scale (*29*). We followed these lines of inquiry by investigating recent TE diversity within each genome of our dataset by calculating the Shannon Diversity Index of TE classes. Focusing on recently inserted TEs, we summed the bases that were attributed to TEs with K2P values less than 4%. We then generated the proportions (*p*_*i*_) for each TE class attributed to the overall base pair total of recently inserted TEs. To calculate the Shannon Diversity Index, *H*, per the equation.

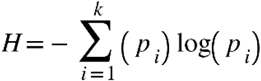

To calculate the evenness of recent TE accumulation among the 5 main categories of TEs, we employed the ecological metric, Pielou’s *J*, a measure of species evenness, where *S* was equal to the total number of recent TE hits found within an assembly.

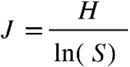

### Dietary data

We gathered diet classification from The Animal Diversity Web (animaldiversity.org) for 178 available mammals on the public database (table S8). The young DNA transposon dataset was then compared against three diet types: carnivore, herbivore, and omnivore.

### Hierarchical Bayesian analyses

A hierarchical Bayesian approach was adopted to simultaneously estimate the species-specific structure of errors while estimating error for the beta-distributed proportion of TE in the genome. A hierarchical approach is often called a mixed model in the literature, with cluster-specific effects called “random”, and sample-wide effects called “fixed”. As different fields apply random and fixed to different levels of the hierarchy, here we adopt the language of cluster-specific and sample-wide effects (*62*). Analyses begin by modelling the proportion of genome as a function of the genome assembly size as a beta-distributed variable (*63*):

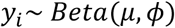

In which *μ* is the mean, and *ϕ* relates to the variance such that:

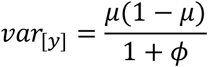

Given observations *Y*, and covariate assembly size *X*:

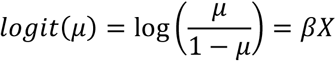

Instead of a typical regression, in which observations are presumed to be independent, our analyses account for the phylogenetic structure of the errors by including normally distributed species-specific effects with phylogenetic errors (*64*) such that:

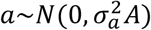

In which the phylogenetic relationship matrix *A* (*65*) replaces the identity of observations for the residuals. The same distribution of the response and its phylogenetic errors was applied across all regressions.

Assembly sizes in base pairs were on the order of 10^9^. To enable efficient modeling, this predictor was log10 transformed and then scaled (subtracting the mean and dividing by one standard deviation). No other predictor variables were transformed. Analyses of the association between diet and TE proportions used diet as a group-specific predictor.

To implement Bayesian sampling for these analyses, we used brms (*66*), a package that enables coding models in R for implementation in the stan statistical language (*67*). We ran separate univariate models for each set of predictors (assembly size, diet, Shannon’s Diversity Index, and Pielou’s Evenness Index), with the proportion of TE in the genome as the response. The covariance matrix A was obtained from the variance covariance matrix of the dated phylogeny (*65*) of sampled species. Models ran four separate Markov chain Monte Carlo chains using a Hamiltonian Monte Carlo approach. Compared to other Bayesian implementations, the HMC approach saves time in sampling parameter spaces by generating efficient transitions spanning the posterior based on derivatives of the density function of the model. We used the approach of (*68*) to estimate R^2^ from hierarchical Bayesian models. This approach divides the variance of the predicted values by the variance of predicted values plus the expected variance of the errors.

## Supporting information

Data Speadsheets

Supplemental Figures & Spreadsheet Data Descriptions

## Acknowledgements

We thank the High-Performance Computing Center at Texas Tech University for providing compute resources and technical support throughout the project. This work was also made possible by the SeaWulf computing system from Stony Brook Research Computing and Cyberinfrastructure, and the Institute for Advanced Computational Science at Stony Brook University funded by National Science Foundation-OAC 1531492. We also thank Brittany Ann Hale for providing artistic renditions of mammal taxa for our figure.

## Funding

This project was partially supported by

National Science Foundation grant DEB 1838283 (DDMS, DAR)

National Science Foundation grant IOS 2032006 (DDMS, DAR)

National Institutes of Health grant R01HG002939 (JMS, RH, AS, JebR)

National Institutes of Health grant U24HG010136 (JMS, RH, AS, JebR)

National Science Foundation grant DEB 1838273 (LMD)

National Science Foundation grant DGE 1633299 (LMD)

National Institutes of Health NHGRI R01HG008742 (ZC)

Swedish Research Council Distinguished Professor Award (ZC)

## Author contributions

Conceptualization: ABO, DAR

Assembly generation: DDMS, LMD, DAR

Library validation & curation: NSP, JMS, ABO, KAMS, JK, JRG, MH, CG, CC, JR, JebR, RH, AS, DAR

Methodology & Investigation: ABO, LMD, NSP, DAR

Writing – original draft: ABO NSP, DAR, RDS, LMD

Writing – review & editing: ABO, NSP, DAR, JMS, AS, RDS, LMD

## Data and materials availability

All assemblies are available in Genbank, TE consensus sequences are available via the Dfam database. All other data is available in the supplementary materials; code used in the analysis is available at (*49*)..

## Zoonomia Consortium List

Gregory Andrews^1^, Joel C. Armstrong^2^, Matteo Bianchi^3^, Bruce W. Birren^4^, Kevin R. Bredemeyer^5^, Ana M. Breit^6^, Matthew J. Christmas^3^, Hiram Clawson^2^, Joana Damas^7^, Federica Di Palma^8,9^, Mark Diekhans^2^, Michael X. Dong^3^, Eduardo Eizirik^10^, Kaili Fan^1^, Cornelia Fanter^11^, Nicole M. Foley^5^, Karin Forsberg-Nilsson^12,13^, Carlos J. Garcia^14^, John Gatesy^15^, Steven Gazal^16^, Diane P. Genereux^4^, Linda Goodman^17^, Jenna Grimshaw^14^, Michaela K. Halsey^14^, Andrew J. Harris^5^, Glenn Hickey^18^, Michael Hiller^19,20,21^, Allyson G. Hindle^11^, Robert M. Hubley^22^, Graham M. Hughes^23^, Jeremy Johnson^4^, David Juan^24^, Irene M. Kaplow^25,26^, Elinor K. Karlsson^1,4,27^, Kathleen C. Keough^17,28,29^, Bogdan Kirilenko^19,20,21^, Klaus-Peter Koepfli^30,31,32^, Jennifer M. Korstian^14^, Amanda Kowalczyk^25,26^, Sergey V. Kozyrev^3^, Alyssa J. Lawler^4,26,33^, Colleen Lawless^23^, Thomas Lehmann^34^, Danielle L. Levesque^6^, Harris A. Lewin^7,35,36^, Xue Li^1,4,37^, Abigail Lind^28,29^, Kerstin Lindblad-Toh^3,4^, Ava Mackay-Smith^38^, Voichita D. Marinescu^3^, Tomas Marques-Bonet^39,40,41,42^, Victor C. Mason^43^, Jennifer R. S. Meadows^3^, Wynn K. Meyer^44^, Jill E. Moore^1^, Lucas R. Moreira^1,4^, Diana D. Moreno-Santillan^14^, Kathleen M. Morrill^1,4,37^, Gerard Muntané^24^, William J. Murphy^5^, Arcadi Navarro^39,41,45,46^, Martin Nweeia^47,48,49,50^, Sylvia Ortmann^51^, Austin Osmanski^14^, Benedict Paten^2^, Nicole S. Paulat^14^, Andreas R. Pfenning^25,26^, BaDoi N. Phan^25,26,52^, Katherine S. Pollard^28,29,53^, Henry E. Pratt^1^, David A. Ray^14^, Steven K. Reilly^38^, Jeb R. Rosen^22^, Irina Ruf^54^, Louise Ryan^23^, Oliver A. Ryder^55,56^, Pardis C. Sabeti^4,57,58^, Daniel E. Schäffer^25^, Aitor Serres^24^, Beth Shapiro^59,60^, Arian F. A. Smit^22^, Mark Springer^61^, Chaitanya Srinivasan^25^, Cynthia Steiner^55^, Jessica M. Storer^22^, Kevin A. M. Sullivan^14^, Patrick F. Sullivan^62,63^, Elisabeth Sundström^3^, Megan A. Supple^59^, Ross Swofford^4^, Joy-El Talbot^64^, Emma Teeling^23^, Jason Turner-Maier^4^, Alejandro Valenzuela^24^, Franziska Wagner^65^, Ola Wallerman^3^, Chao Wang^3^, Juehan Wang^16^, Zhiping Weng^1^, Aryn P. Wilder^55^, Morgan E. Wirthlin^25,26,66^, James R. Xue^4,57^, Xiaomeng Zhang^4,25,26^

Affiliations:

^1^Program in Bioinformatics and Integrative Biology, UMass Chan Medical School; Worcester, MA 01605, USA.

^2^Genomics Institute, University of California Santa Cruz; Santa Cruz, CA 95064, USA.

^3^Department of Medical Biochemistry and Microbiology, Science for Life Laboratory, Uppsala University; Uppsala, 751 32, Sweden.

^4^Broad Institute of MIT and Harvard; Cambridge, MA 02139, USA.

^5^Veterinary Integrative Biosciences, Texas A&M University; College Station, TX 77843, USA.

^6^School of Biology and Ecology, University of Maine; Orono, ME 04469, USA. ^7^The Genome Center, University of California Davis; Davis, CA 95616, USA.

^8^Genome British Columbia; Vancouver, BC, Canada.

^9^School of Biological Sciences, University of East Anglia; Norwich, UK.

^10^School of Health and Life Sciences, Pontifical Catholic University of Rio Grande do Sul; Porto Alegre, 90619-900, Brazil.

^11^School of Life Sciences, University of Nevada Las Vegas; Las Vegas, NV 89154, USA.

^12^Biodiscovery Institute, University of Nottingham; Nottingham, UK.

^13^Department of Immunology, Genetics and Pathology, Science for Life Laboratory, Uppsala University; Uppsala, 751 85, Sweden.

^14^Department of Biological Sciences, Texas Tech University; Lubbock, TX 79409, USA.

^15^Division of Vertebrate Zoology, American Museum of Natural History; New York, NY 10024, USA.

^16^Keck School of Medicine, University of Southern California; Los Angeles, CA 90033, USA.

^17^Fauna Bio Incorporated; Emeryville, CA 94608, USA.

^18^Baskin School of Engineering, University of California Santa Cruz; Santa Cruz, CA 95064, USA.

^19^Faculty of Biosciences, Goethe-University; 60438 Frankfurt, Germany.

^20^LOEWE Centre for Translational Biodiversity Genomics; 60325 Frankfurt, Germany.

^21^Senckenberg Research Institute; 60325 Frankfurt, Germany.

^22^Institute for Systems Biology; Seattle, WA 98109, USA.

^23^School of Biology and Environmental Science, University College Dublin; Belfield, Dublin 4, Ireland.

^24^Department of Experimental and Health Sciences, Institute of Evolutionary Biology (UPF-CSIC), Universitat Pompeu Fabra; Barcelona, 08003, Spain.

^25^Department of Computational Biology, School of Computer Science, Carnegie Mellon University; Pittsburgh, PA 15213, USA.

^26^Neuroscience Institute, Carnegie Mellon University; Pittsburgh, PA 15213, USA.

^27^Program in Molecular Medicine, UMass Chan Medical School; Worcester, MA 01605, USA.

^28^Department of Epidemiology & Biostatistics, University of California San Francisco; San Francisco, CA 94158, USA.

^29^Gladstone Institutes; San Francisco, CA 94158, USA.

^30^Center for Species Survival, Smithsonian’s National Zoo and Conservation Biology Institute; Washington, DC 20008, USA.

^31^Computer Technologies Laboratory, ITMO University; St. Petersburg 197101, Russia.

^32^Smithsonian-Mason School of Conservation, George Mason University; Front Royal, VA 22630, USA.

^33^Department of Biological Sciences, Mellon College of Science, Carnegie Mellon University; Pittsburgh, PA 15213, USA.

^34^Senckenberg Research Institute and Natural History Museum Frankfurt; 60325 Frankfurt am Main, Germany.

^35^Department of Evolution and Ecology, University of California Davis; Davis, CA 95616, USA.

^36^John Muir Institute for the Environment, University of California Davis; Davis, CA 95616, USA.

^37^Morningside Graduate School of Biomedical Sciences, UMass Chan Medical School; Worcester, MA 01605, USA.

^38^Department of Genetics, Yale School of Medicine; New Haven, CT 06510, USA.

^39^Catalan Institution of Research and Advanced Studies (ICREA); Barcelona, 08010, Spain.

^40^CNAG-CRG, Centre for Genomic Regulation, Barcelona Institute of Science and Technology (BIST); Barcelona, 08036, Spain.

^41^Department of Medicine and LIfe Sciences, Institute of Evolutionary Biology (UPF-CSIC), Universitat Pompeu Fabra; Barcelona, 08003, Spain.

^42^Institut Català de Paleontologia Miquel Crusafont, Universitat Autònoma de Barcelona; 08193, Cerdanyola del Vallès, Barcelona, Spain.

^43^Institute of Cell Biology, University of Bern; 3012, Bern, Switzerland.

^44^Department of Biological Sciences, Lehigh University; Bethlehem, PA 18015, USA.

^45^BarcelonaBeta Brain Research Center, Pasqual Maragall Foundation; Barcelona, 08005, Spain.

^46^CRG, Centre for Genomic Regulation, Barcelona Institute of Science and Technology (BIST); Barcelona, 08003, Spain.

^47^Department of Comprehensive Care, School of Dental Medicine, Case Western Reserve University; Cleveland, OH 44106, USA.

^48^Department of Vertebrate Zoology, Canadian Museum of Nature; Ottawa, Ontario K2P 2R1, Canada.

^49^Department of Vertebrate Zoology, Smithsonian Institution; Washington, DC 20002, USA.

^50^Narwhal Genome Initiative, Department of Restorative Dentistry and Biomaterials Sciences, Harvard School of Dental Medicine; Boston, MA 02115, USA.

^51^Department of Evolutionary Ecology, Leibniz Institute for Zoo and Wildlife Research; 10315 Berlin, Germany.

^52^Medical Scientist Training Program, University of Pittsburgh School of Medicine; Pittsburgh, PA 15261, USA.

^53^Chan Zuckerberg Biohub; San Francisco, CA 94158, USA.

^54^Division of Messel Research and Mammalogy, Senckenberg Research Institute and Natural History Museum Frankfurt; 60325 Frankfurt am Main, Germany.

^55^Conservation Genetics, San Diego Zoo Wildlife Alliance; Escondido, CA 92027, USA.

^56^Department of Evolution, Behavior and Ecology, School of Biological Sciences, University of California San Diego; La Jolla, CA 92039, USA.

^57^Department of Organismic and Evolutionary Biology, Harvard University; Cambridge, MA 02138, USA.

^58^Howard Hughes Medical Institute; Chevy Chase, MD, USA.

^59^Department of Ecology and Evolutionary Biology, University of California Santa Cruz; Santa Cruz, CA 95064, USA.

^60^Howard Hughes Medical Institute, University of California Santa Cruz; Santa Cruz, CA 95064, USA.

^61^Department of Evolution, Ecology and Organismal Biology, University of California Riverside; Riverside, CA 92521, USA.

^62^Department of Genetics, University of North Carolina Medical School; Chapel Hill, NC 27599, USA.

^63^Department of Medical Epidemiology and Biostatistics, Karolinska Institutet; Stockholm, Sweden.

^64^Iris Data Solutions, LLC; Orono, ME 04473, USA.

^65^Museum of Zoology, Senckenberg Natural History Collections Dresden; 01109 Dresden, Germany.

^66^Allen Institute for Brain Science; Seattle, WA 98109, USA

## Supplementary Materials

Figs. S1-S5

Tables S1-S8

