## Supplemental Figures & Spreadsheet Data Descriptions for "Insights into mammalian TE diversity via the curation of 248 mammalian genome assemblies"

Science
AAAS

### Supplementary Materials for

#### Insights into mammalian TE diversity via the curation of 248 mammalian genome assemblies

Austin B. Osmanski[1], Nicole S. Paulat[1], Jenny Korstian[1], Jenna R. Grimshaw[1], Michaela Halsey[1], Kevin A.M. Sullivan[1], Diana D. Moreno-Santillán[1], Claudia Crookshanks[1], Jacquelyn Roberts[1], Carlos Garcia[1], Liliana M. Dávalos[4,5], Matthew G. Johnson[1], Llewellyn D. Densmore[1], Richard D. Stevens[2], Zoonomia Consortium, Jeb Rosen[3], Jessica M. Storer[3], Robert Hubley[3], Arian F.A. Smit[3], David A. Ray[1*]

**This PDF file includes:**

Figs. S1-S5
Tables S1-S8

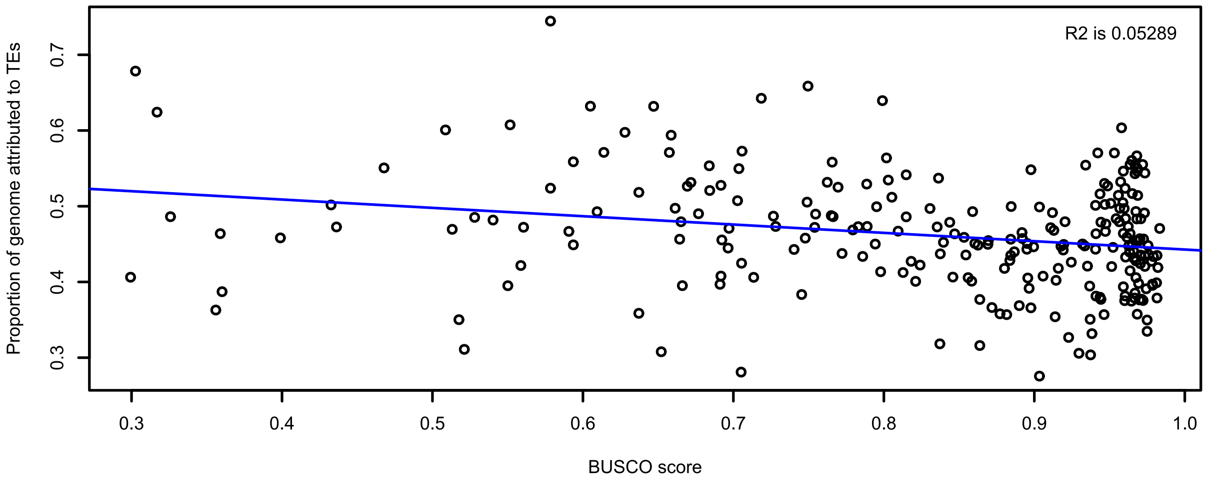


**Fig. S1.**

BUSCO quality metric compared with overall proportion of genomic content attributed to TEs for each mammal assembly.

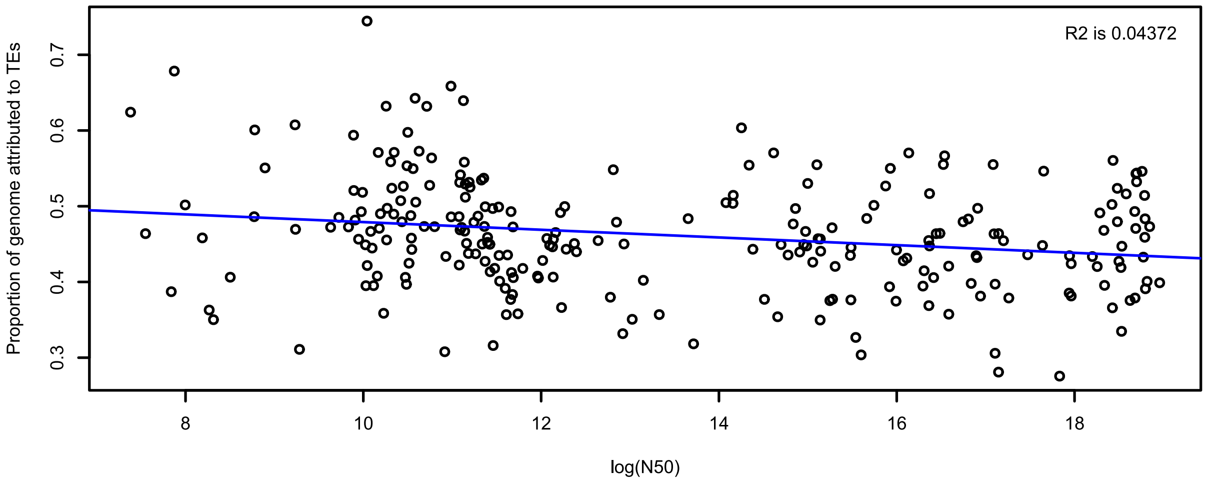


**Fig. S2.**

N50 quality metric compared with overall proportion of genomic content attributed to TEs for each mammal assembly.

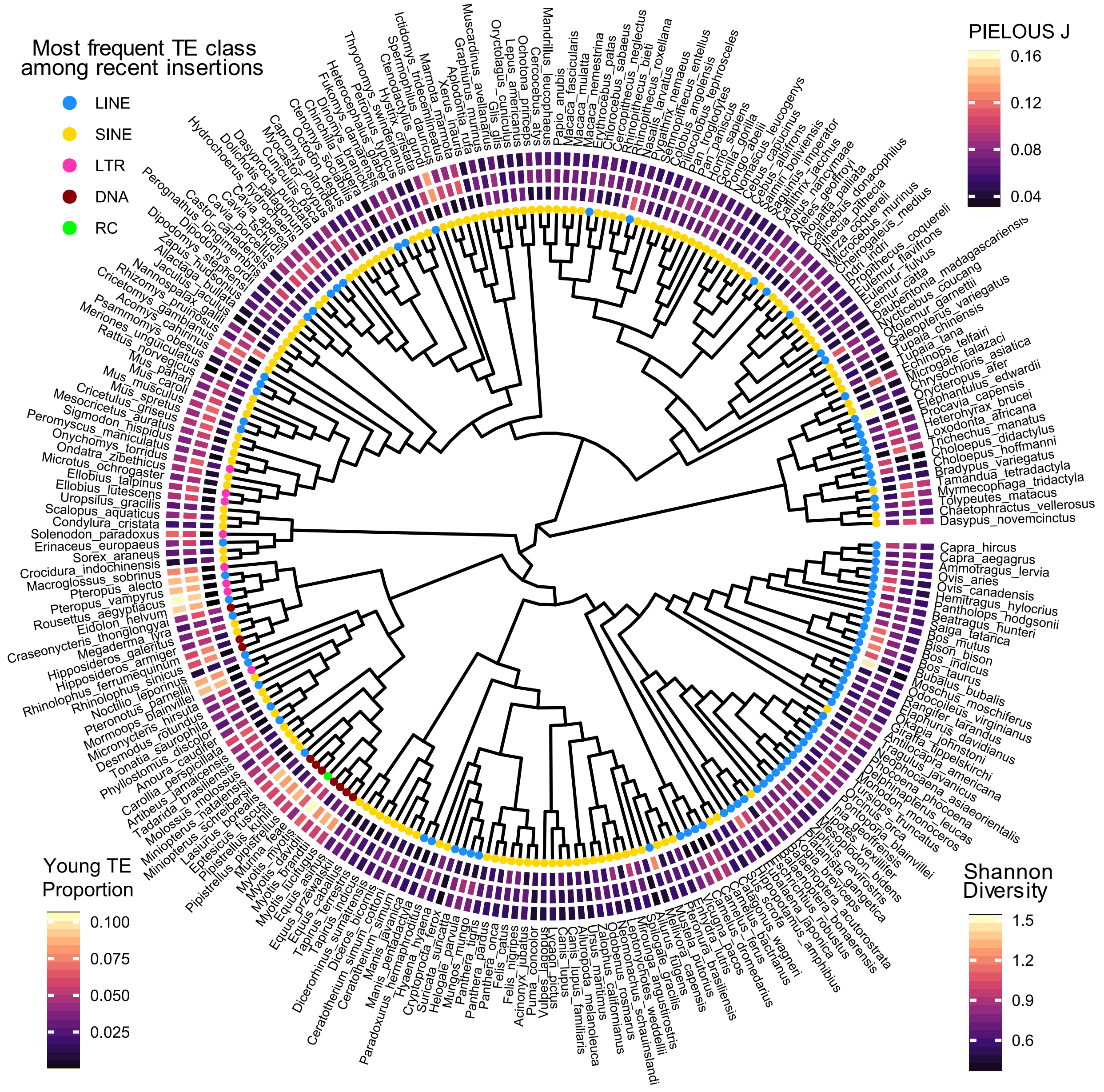


**Fig. S3.**

TE diversity in mammals within a phylogenetic context. Circles at tips of the tree represent the most common TE class among recent insertions. The inner ring depicts a heatmap representing the proportion of recent TE insertions within each assembly. The middle ring signifies the

Shannon Diversity Index, with higher values indicating a more diverse TE landscape within a given species. The outer ring depcits a measure of TE accumulation evenness as calculated by Pielous J. Phylogeny adapted from (*64*).

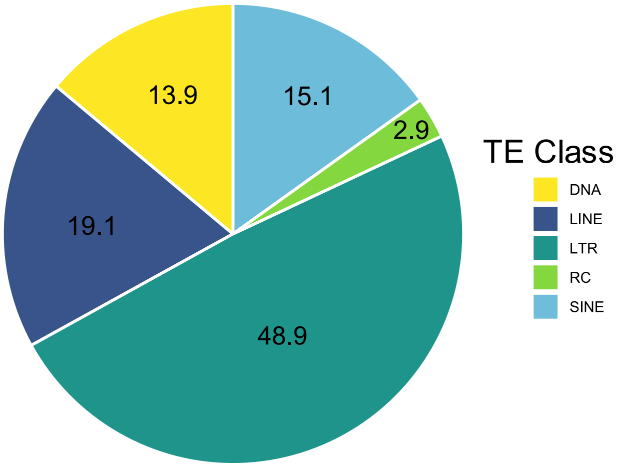


**Fig. S4.**

This figure illustrates the proportion of each TE class among all novel elements (n=8,258) curated during this project. Novel elements are TE sequences we curated but had yet to be included within the Repbase (*2*) database before the initiation of this project.

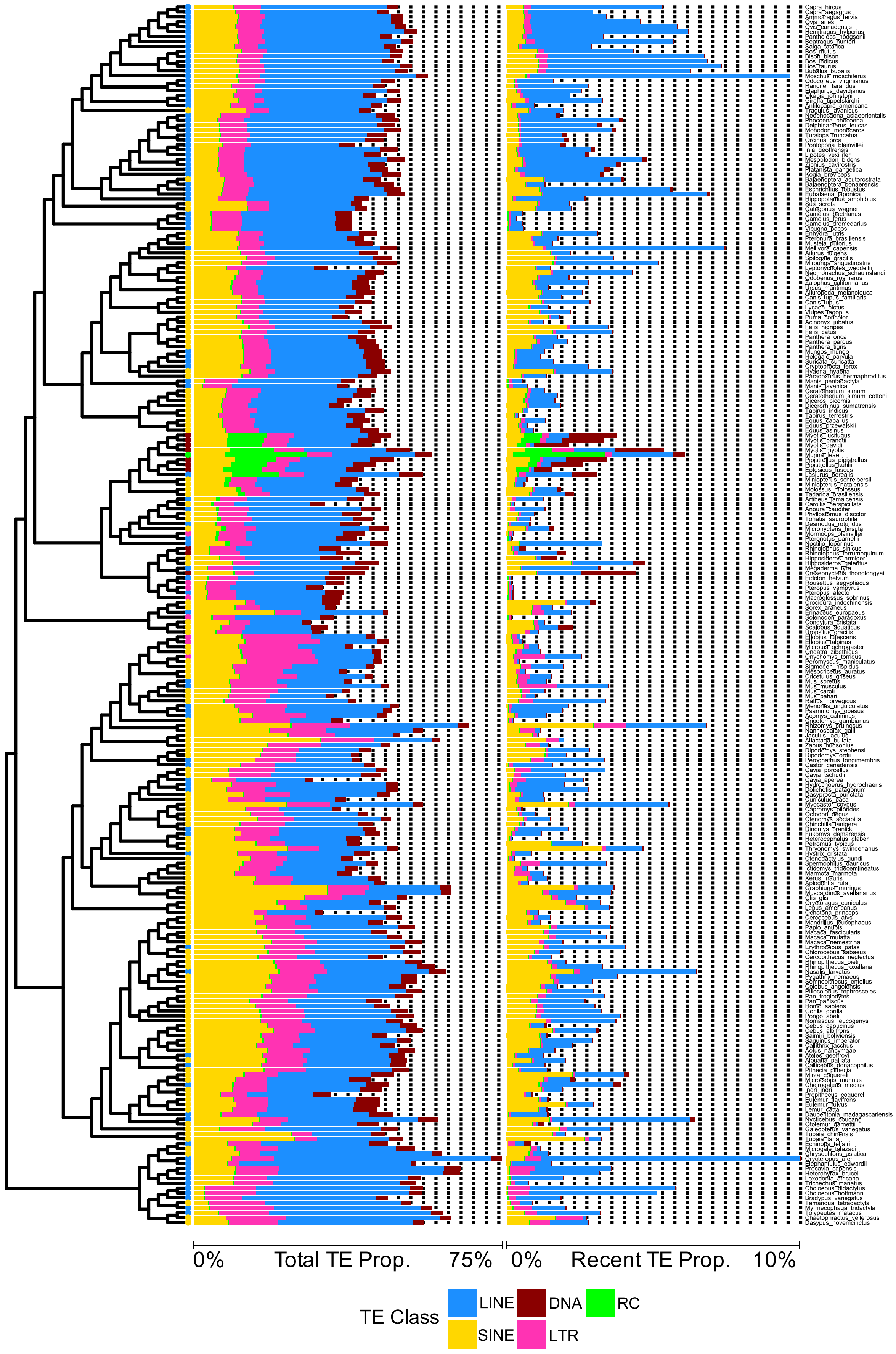
0%
Total TE Prop.
75%
0%
Recent TE Prop.
10%
TE Class
LINE
DNA
RC
SINE
LTR

**Fig. S5.**

This phylogeny is a linear representation of the circular version found within the main manuscript (Fig 2).

**Table S1.**

Spreadsheet tab label: “1 AssemblySize_v_TEprop.” Summary of posterior distributions of the coefficient of proportion of genome by TEs as a function of assembly size. Posterior distributions excluding 0 are in bold, indicating significance. ESS, Estimated sampling size; HPD, high probability density interval; l, lower; PSRF, potential scale reduction factor; u, upper. Note that parameters of the beta regression define change in the logit scale and thus need to be exponentiated to be interpreted as change in units.

**Table S2.**

Spreadsheet tab label: “S2 Species_List.” List of 248 placental mammal species used for TE annotation. Species indicated with an asterisk were obtained from the bat1K project, all others are from the Zoonomia Consortium. Subcategories indicate species that were excluded from TE library curation analyses.

**Table S3.**

Spreadsheet tab label: “S3 TE_proportions.” TE proportions and genome size (base pairs) for each species.

**Table S4.**

Spreadsheet tab label: “S4 Recent_TE_proportions.” TE proportions of recently inserted TEs (kimura < 4.4) and genome size (base pairs) for each species.

**Table S5.**

Spreadsheet tab label: "S5 Diversity." Measures of TE diversity, Shannon *H* and Pielou's *J*, per species.

**Table S6.**

Spreadsheet tab label: “S6 Shannon_v_YoungTE.” Summary of posterior distributions of the coefficient of proportion of genome by recently accumulated TEs as a function of Shannon’s Diversity Index.  ESS, Estimated sampling size; HPD, high probability density interval; l, lower; PSRF, potential scale reduction factor; u, upper. Note that parameters of the beta regression define change in the logit scale and thus need to be exponentiated to be interpreted as change in units, and thus define a curvilinear relationship.

**Table S7.**

Spreadsheet tab label: "S7 Pielous_v_YoungTE." Summary of posterior distributions of the coefficient of proportion of genome by recently accumulated TEs as a function of Pielou's Diversity Index. ESS, Estimated sampling size; HPD, high probability density interval; l, lower; PSRF, potential scale reduction factor; u, upper. Note that parameters of the beta regression define change in the logit scale and thus need to be exponentiated to be interpreted as change in units, and thus define a curvilinear relationship.

**Table S8.**

Spreadsheet tab label: "S8 Diet." Dietary habit data acquired from Animal Diversity Web (animaldiversity.org).
